# The serotonin transporter genotype x cumulative early-life stress interaction has opposite effects on anxiety-like behaviors in male and female rats

**DOI:** 10.64898/2026.09.04.749397

**Authors:** Mayerli A. Prado-Rivera, Annemarijn M.J. Fortuin, Cato M. Drion, Jocelien D.A. Olivier

## Abstract

Early-life psychosocial adversity increases vulnerability to stress-related disorders and may begin as early as prior to conception via maternal stress. Consequently, early-life stress (ELS) in offspring may accumulate across early development. The short-allele variant of the serotonin transporter (SERT) gene has been implicated in depression risk and higher anxiety sensitivity following stress exposure, but for *cumulative* ELS it is unknown whether behavioral effects depend on the SERT genotype. In this study, we used heterozygous SERT knockout (SERT^+/-^) rats to model reduced SERT expression similar to human short-allele carriers. To investigate the effects of cumulative ELS on offspring behavioral phenotype, dams were exposed to stress before, during and after pregnancy, combined with offspring postnatal maternal separation. Cognitive, affective, and social behaviors were assessed in wild-type (WT) and SERT^+/-^ (HET) male and female adults. Maternal care and anhedonia-like behaviors in dams were also assessed. Dams exposed to the cumulative stress (STR) displayed increased arched-back licking and grooming compared to controls (CTR). In the offspring, HET rats showed increased social investigation and tended to display hyperactivity compared to WT. Anxiety-like behavior was higher in HET females compared to WT females, with no genotype differences in males. Independent of genotype, STR rats consumed less sucrose and showed impaired object recognition compared to CTR; however, HET-STR rats performed better in the object recognition than WT-STR rats. A significant SERT genotype x cumulative ELS x sex interaction revealed opposing, sex-dependent effects on anxiety-like behavior. HET-STR females showed more anxiety-like responses relative to HET-CTR and WT-STR females, as well as HET-STR males. In contrast, HET-STR males were less anxious than HET-CTR and WT-STR males. Reduced SERT expression is linked to more social investigation alongside increased anxiety, particularly in SERT^+/-^ females. While cumulative ELS did not induce robust anxiety-like and depressive-like phenotypes, overall, it impaired memory performance, an effect partially mitigated in SERT^+/-^ rats. Notably, the SERT genotype differentially modulated anxiety-like behavior in male versus female offspring, increasing vulnerability in SERT^+/-^ females while reducing it in SERT^+/-^ males, an effect further amplified by exposure to cumulative ELS. These results underscore sex as a critical determinant in the interaction between genetic susceptibility and cumulative exposure to early adversity in shaping affective outcomes.

## 1. Introduction

Early-life psychosocial adversity has a large impact on mental health and increases risk for psychopathology later in life. Physical and emotional childhood maltreatment is linked to cognitive impairments and poorer interpersonal relations, as well as to higher risk for anxiety disorders and major depression (Bader & Frank, 2024; Danese & Widom, 2024; Kim et al., 2024; Li et al., 2016; Qian et al., 2022; Struck et al., 2021). Psychosocial adversity during pregnancy (e.g., bereavement, domestic violence) has also been associated with negative impact on children’s development of cognition, behavior, and affect (Henrichs et al., 2011; King et al., 2012; Moss et al., 2017; van Den Bergh et al., 2020). In addition, maternal exposure to major life stressors, violence, and war prior to conception has been linked to greater risk of behavioral and cognitive difficulties as well as depression and mood disorders in offspring (Bowers & Yehuda, 2016; Roth et al., 2014; Shachar-Dadon et al., 2017; Yehuda et al., 2008). When such adversity results in cumulative stress over time, as is often the case in low-income households (Houweling & Grünberger, 2024), offspring may consequently experience *cumulative* early-life stress (ELS) exposure across the preconception, prenatal, and postnatal periods.

Offspring genetic makeup interacting with ELS to influence the onset of stress-related disorders has been actively investigated. The serotonin-transporter-linked promoter region (5-HTTLPR) contains a repetitive GC-rich sequence of 20-23-bp-long repeat elements in the gene’s upstream regulatory region, with insertion/deletion variants producing the 14-repeat short (S, low expressing) and 16-repeat long (L, high expressing) alleles. It has been shown that S-allele carriers are more susceptible to depression following exposure to childhood maltreatment, abuse, neglect or chronic family stress than L-allele carriers (Caspi et al., 2003; Jenness M.A. et al., 2011; Y.-K. Kim et al., 2019; Vrijsen et al., 2015). Similarly, S-allele carriers exhibit heightened anxiety sensitivity and an elevated risk of posttraumatic stress disorders following exposure to childhood emotional and physical maltreatment (Cicchetti et al., 2007; Stein et al., 2008; Xie et al., 2009). The role of 5-HTTLPR in psychopathology remains controversial however, as susceptibility of stress-exposed S-allele carriers to depressive and anxiety disorders has been not always found (Border et al., 2019; Bosker et al., 2010; Culverhouse et al., 2018; Özçürümez et al., 2018; Schinka et al., 2004). One reason for this may be that sustained maternal stress, beginning before child’s conception, has not been taken into account.

To our knowledge, no one has studied whether anxiety or depressive symptoms in adult offspring is mediated by the interaction of *cumulative* ELS exposure and the SERT genotype. To study this in a controlled environment, heterozygous SERT knockout (SERT^+/-^) rodents provide a valid model. Although lacking the human 5-HTTLPR, SERT^+/-^ rodents show neurochemical similarities to human S-allele carriers (Bengel et al., 1998; Homberg et al., 2007). Exposure to pregestational, prenatal or postnatal stress increased some depressive-like responses in SERT^+/-^ mouse and rat offspring (Houwing, Ramsteijn, et al., 2019; Houwing, Schuttel, et al., 2020; van Den Hove et al., 2011; van der Doelen et al., 2013), but other studies have found no increased anxiety-like behaviors, or findings appear to depend on sex and ELS timing (Jones et al., 2010; Matsui et al., 2018; Woo et al., 2023). Importantly, such studies investigated individual ELS exposures and did not examine the effects of cumulative ELS on the offspring. The goal of the present study was to examine whether male and female SERT^+/-^ offspring were more vulnerable to developing anxiety-like and depressive-like behaviors following *cumulative* ELS. To do so, we assessed affective, cognitive, and social behaviors in adult wildtype and SERT^+/-^ offspring exposed to maternal stress before, during, and after pregnancy, combined with direct postnatal stress. In addition, maternal behavior and anhedonia were assessed in dams. We hypothesized that: 1) cumulative ELS would increase offspring vulnerability to anxiety-like and depressive-like behaviors and 2) SERT^+/-^ rats would show more pronounced maladaptive behaviors than wildtypes following cumulative ELS.

## 2. Methods

### 2.1. Animals

Twenty-three female Wistar rats (>10 weeks-old; weight: 189.78 ± 9.57 g) from Charles River Laboratories (The Netherlands) and their offspring were socially housed (2-4 animals per cage), unless stated otherwise, under standard laboratory conditions with a reversed 12h:12h light/dark cycle (lights off at 8:00 AM), and 40-60% room humidity. During pregnancy and lactation, dams were individually housed in Makrolon type 3 cages (38.2×22.0×15.0 cm) and nesting material was provided (Enviro-dri®). During social housing, animals were housed in Makrolon type 4 cages (55.6×33.4×19.5 cm), provided with wooden gnawing sticks (10×2×2 cm) and polycarbonate small boxes for enrichment. Access to food (RMH-B, AB Diets; Woerden, the Netherlands) and tap water was *ad libitum*, unless stated otherwise. All breeding occurred in our facilities. Behavioral testing took place between 9:00 AM and 6:00 PM. All experimental procedures were approved by the Institutional Animal Care and Use Committee of The University of Groningen and were conducted in full compliance with the EU Directive (2010/63/EU).

### 2.2. Experimental design

Females were randomized to control (CTR, n=10) or cumulative stress (STR, n= 13). For STR offspring to be exposed to cumulative ELS, no cross-fostering was performed. STR females underwent chronic unpredictable stress (CUS) immediately *before* pregnancy (pregestational), unpredictable restraint stress (URS) *during* the last week of pregnancy (prenatal), and restraint stress or forced swimming *after* pregnancy (postnatal) while pups were exposed to unpredictable maternal separation (UMS) during the first two postnatal weeks (for detailed pregestational, prenatal and postnatal stress protocols see Suppl. File 1). CTR females were left undisturbed. Maternal behaviors were assessed from postnatal day (P)1 to P7. Pups were weaned on P21, anhedonia in dams was assessed 2±1 weeks after, and were euthanized afterwards. Offspring were behaviorally tested between P90(±10) and P160(±10) and then euthanized.

**Fig1.**
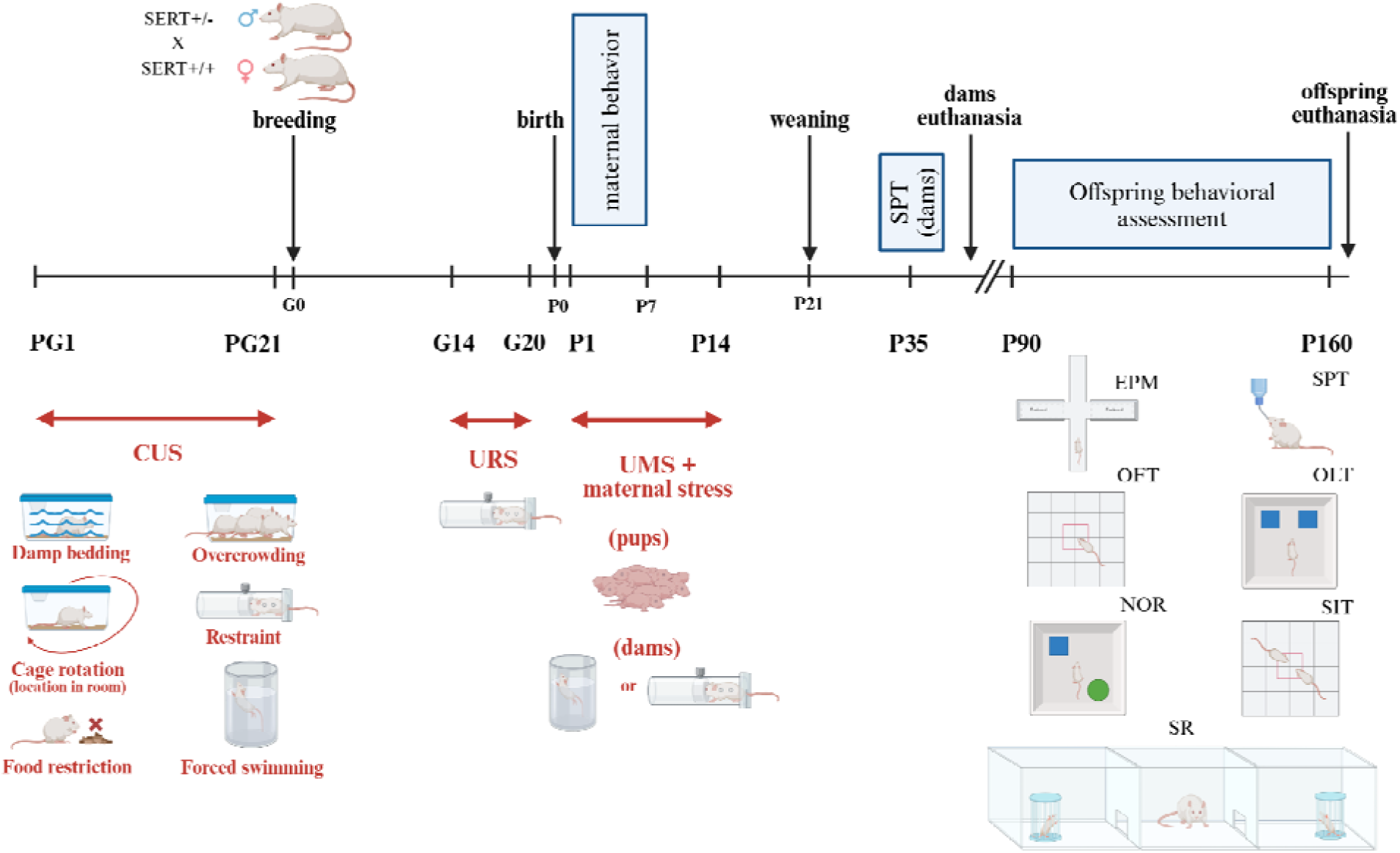
Schematic representation of experimental design. Offspring exposure to *cumulative ELS* across the preconception, prenatal, and postnatal periods. Dams were daily exposed to CUS for three weeks immediately before breeding and URS during the last week of pregnancy. Offspring were exposed to UMS for 3 hr/day while dams underwent restraint stress or forced swimming. CUS: Chronic unpredictable stress; URS: Unpredictable restraint stress; UMS: Unpredictable maternal separation. PG: Pregestational day; G: Gestational day; P: Postnatal day. SPT: Sucrose preference test; EPM: Elevated plus maze; OFT: Open field test; OLT: Object location test; NOR: Novel object recognition test; SIT: Social interaction test; SR: Social recognition test.

### 2.3. Behavioral testing in dams

#### 2.3.1. Maternal care assessment

CTR- and STR-dams were observed three times daily from P1-P7. All observation sessions occurred during the dark part of the reverse light/dark cycle, before UMS (session 1), immediately after UMS (session 2), and 2-3 hr after UMS (session 3). Each session consisted of 30 min, during which dam’s frequency on the nest and activity were scored once per minute (resulting in 30 1-min epochs). Dam’s frequency on the nest was scored when the dam was within the nest she built with nesting material. Activities scored were pup-directed behaviors (arched-back nursing (AB), licking/grooming (LG), nest building, and pup retrieval), and self-directed behaviors (drinking, eating, self-grooming, and resting).

For each session, dam’s frequency on the nest was calculated as the number of epochs the dam was observed on the nest, divided by the total number of epochs (30), multiplied by 100. The percentage of frequency of dams engaged in pup-directed behaviors, and dams engaged in mixed care (i.e., displaying both pup- and self-directed behaviors) were also calculated. To further explore active maternal care, the percentage of frequency of dams displaying AB/LG was also calculated. P1 to P7 were grouped to get the total percentage of frequency for each one of these outcomes per session.

#### 2.3.2. Sucrose preference test (SPT)

Two±1 weeks after weaning the litter, dams were habituated to two bottles of water, one on each side of the cage. 24 hr later, they were exposed to one bottle of water and one bottle containing a sucrose solution for 24 hr on alternating days. On the other days two bottles of water were presented. With each sucrose day, the sucrose concentration increased (0.5%, 1%, 2%, 4%). Sucrose bottle locations on the cage were alternated on sucrose days to prevent spatial bias. Fluid consumption (gram) was determined daily by comparing bottles weight before and after consumption. The preference for sucrose over water was calculated as:

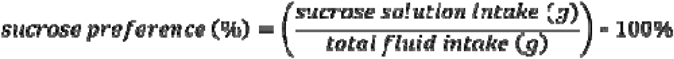

Actual sucrose intake in mg per gram of rat weight -corrected for body weight-was also calculated as:

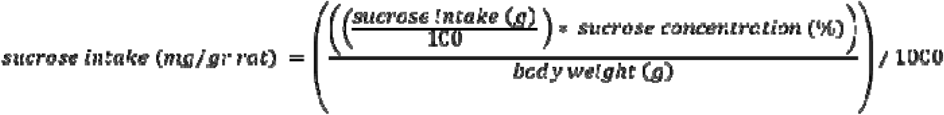

The day after completion of the SPT, females were euthanized by induction of deep anesthesia with isoflurane followed by rapid decapitation, and brain tissue was collected for analysis in a separate study.

### 2.4. Behavioral testing in adult offspring

Pups were weaned on P21 and housed with their litter mates until P30. Then, 2-4 rats of both genotypes were caged with mates of same sex and ELS for the rest of the study. To minimize litter effects, no more than one pup per genotype and sex was included from each litter whenever feasible. Behavioral testing was conducted between P90(±10) and P160(±10) in eight groups: WT-CTR, HET-CTR, WT-STR, and HET-STR for both males and females (n=10 per genotype/sex/ELS). Tests were conducted at least one week apart in the following order: elevated plus maze (EPM), sucrose preference test (SPT), open field test (OFT), object location test (OLT), novel object recognition (NOR), social interaction test (SIT), and social recognition (SR) (for detailed tasks procedures see Suppl. File 1). Approximately 1 week after completion of the last behavioral test, rats were euthanized by induction of deep anesthesia with isoflurane followed by rapid decapitation, and brain tissue was collected for analysis in a separate study.

### 2.5. Statistical analysis

All statistical analyses were carried out in R version 4.4.3 (R Core Team, 2024). The statistically significant level was set as p<0.05.

#### 2.8.1. Analysis of dams’ outcomes

Using daily body weight data (Suppl. Table S2A-C), dams’ weight gain per ELS timing (i.e., pregestational, prenatal, and postnatal) was analyzed. Baseline measurement per ELS timing was set as follows: day of arrival as pregestational day (PG) 0; day of breeding as gestational day (G)0; day of delivery as P0. Weight gain per ELS timing was assessed by performing a 2-way mixed ANOVA (day x stress), with day as the within-subjects factor. Data were assessed to identify extreme outliers using the Tukey’s IQR method with the `identify_outliers()` function [rstatix package]. Before running ANOVA, a mixed-linear model was built using `lmer()` function [lme4 package]. Assumptions of normality and homogeneity of variance were checked, and the ANOVA was performed using `Anova()` function [car package]. Post hoc comparisons for repeated-measures factor (day) were corrected using FDR due to the large number of levels. For the between-subject factor (ELS), LSD test was used. When non-normal distribution of residuals was found, raw data was transformed depending on the skewness degree (mild, moderate or severe), the model updated, and the 2-way mixed ANOVA performed. If, even after data transformation the assumptions were not met, an aligned rank transform analysis was chosen for non-parametric testing. In such instances, a mixed-linear model was built using `art()` function [ARTool package], and the ANOVA was performed using `anova()` function [ARTool package]. Maternal care outcomes were analyzed by performing 2-way mixed ANOVAs (session x stress), with session as the within-subjects factor, following the procedures indicated above. For the SPT outcomes, each sucrose concentration was assessed separately, so that independent Welch two sample t-tests were used for CTR vs STR comparisons.

#### 2.8.2. Analysis of offspring’ outcomes

Litters’ weight was assessed via 2-way mixed ANOVA (day x stress), with day as the within-subjects factor. For adult offspring weight, first a 4-way mixed ANOVA (genotype x stress x sex x week-as repeated factor) was performed to determine main sex and/or genotype effects. As a main effect of sex was found but not a main effect of genotype (Suppl. Table S3), 3-way mixed ANOVAs (genotype x stress x week-as repeated factor) were performed separately by sex. Offspring behavioral data were analyzed using 3-way ANOVA (genotype x stress x sex). For SPT outcomes, each sucrose concentration was assessed separately. All 3 factor ANOVAs were followed by 2-way interactions and simple main effects when appropriate. For all offspring outcomes, the same statistical procedures as described for dams were followed.

For the OLT and the NOR, we tested whether each group’s DI was significantly above chance (for detailed DI calculation see Suppl. File 1). Following Akkermand’s et al. (2012) statistical recommendation to reduce Type I error, we generated fictional comparison groups for each experimental group and task, by setting the mean to 0. The variance of each fictional group was estimated from obj1 exploration times during the training trial of the corresponding experimental group, resulting in sixteen fictional comparison groups in total. For this estimation, we calculated the DI of obj1 (DI_obj1_) per animal and these DI_obj1_ values were then averaged to obtain the standard deviation and the standard error of the mean for the corresponding DI_fic_.

## 3. Results

No alteration in general health status of females emerged at the end of the cumulative ELS procedure. Health measures were taken by the researcher through daily observation of animals in their home cage while assessing behavioral and physical indicators of welfare. These included hunched posture, dull or sluggish movements, excessive grooming, absence of feces, rough hair coat, squinted eyes and skin abrasions/lesions (Burkholder et al., 2012).

### 3.1. Dams’ outcomes

#### 3.1.1. Pregnancy outcomes

Pregnancy outcomes of CTR and STR dams are presented in Suppl. Table S4. Eight out of 23 females (3 CTR; 5 STR) did not get pregnant, so a second attempt of breeding was made, during which one STR female failed to maintain her pregnancy. Therefore, a total of 10 CTR and 12 STR females gave birth. Gestational length was not significantly different between CTR and STR dams (t=_(9)_1.50, p=0.17). While pup mortality in the STR group was about twice as low as in the CTR group (7.82% vs. 14.85%), this difference was not statistically significant (OR=0.49, p=0.13). The litters were not culled, and no significant difference in litter size (t_(19.44)_=0.36, p=0.72) or sex ratio (t_(17.29)_=1.25, p=0.23) was found between CTR and STR groups. One CTR female and her litter were excluded from further analysis due to a litter size of three pups, of which only one survived; therefore, further data analysis was conducted for 21 dams and their litters (9 CTR and 12 STR).

#### 3.1.2. Cumulative stress effects on dams’ weight gain

During pregestational stress, females were daily exposed to varied, unpredictable stressors during 21 days immediately before breeding. The weight gain during this period is presented in Suppl. FigS1A. A main effect of day (F_(21,399)_=32.21, p<0.001) and a day x stress interaction effect (F_(21,399)_=5.68, p<0.001) were found. Further analysis revealed that STR females gained significantly less weight than CTR females on PG1 (p<0.05), PG5 (p<0.001), and PG8 (p<0.01); a trend towards a lower weight gain in STR females was observed on PG12 (p=0.06), PG14 (p=0.05), and PG16 (p=0.06). No main exposure effect was found (F_(1,19)_=2.54, p=0.13).

During gestation, body weight was measured daily, while prenatal stress was applied from G14 to G20. Weight gain during this period is presented in Suppl. FigS1B. A main effect of day (F_(20,380)_=600.09, p<0.001), and a day x stress interaction effect (F_(20,380)_=5.71, p<0.001) were observed. Further analysis indicated that, before prenatal intervention, STR dams gained significantly more weight than CTR dams on G12 (p<0.05) and showed a trend towards greater weight gain on G13 (p=0.06) and G14 (p=0.09). Once the prenatal stress started, STR dams progressively gained less weight than CTR dams; a trend towards lower weight gain was found for G17 (p=0.09) and G18 (p=0.09), and the difference was statistically significant on G19 (p<0.05) and G20 (p<0.05). No main effect of stress was found (F_(1,19)_=0.20, p=0.66).

After offspring birth, dams were exposed to stress from P1 to P14 during the period in which UMS was applied to their pups. A main effect of day on dams’ weight gain was observed (F_(15,285)_=64.05, p<0.001), with further analysis indicating a progressive increase over time. No main effect of stress (F_(1,19)_=0.03, p=0.85), or a day x stress interaction effect (F_(15,285)_=0.77, p=0.71) were observed (Suppl. FigS1C).

#### 3.1.3. Cumulative stress effect on dams’ maternal care

One STR dam was excluded from these analyses due to the small litter size (3 pups), which may have affected maternal behaviors; thus, analyses included 9 CTR and 11 STR dams. Maternal care outcomes are presented for session 1 (before UMS), session 2 (immediately after UMS), and session 3 (2-3 hr after UMS), with data pooled across P1-P7 for each session. A main effect of session on the dams’ frequency on the nest was found (F_(2,36)_=57.24, p<0.001; Fig2A), with further analysis revealing a higher frequency in sessions 2 (p<0.001) and 3 (p<0.001) compared to session 1. A main effect of stress was also observed (F_(1,18)_=8.17, p<0.05) where STR dams exhibited a higher percentage in comparison to CTR dams. No session x stress interaction effect (F_(2,36)_=1.07, p=0.35) was found. A main effect of session on pup-directed behaviors was found (F_(2,36)_=55.21, p<0.001; Fig2B), in which pup-directed behaviors were significantly higher in sessions 2 (p<0.001) and 3 (p<0.001) compared to session 1. A main effect of stress was also found (F_(1,18)_=10.33, p<0.01), where STR dams exhibited more pup-directed behaviors than CTR dams. No session x stress interaction effect was found (F_(2,36)_=0.69, p=0.51). No difference in mixed-care behaviors were observed by session (F_(2,36)_=0.06, p=0.95), stress (F_(1,18)_=0.03, p=0.86), or their interaction (F_(2,36)_=1.17, p=0.32) (data not shown). A main effect of session on AB/LG behaviors were observed (F_(2,36)_= 7.48, p<0.01; Fig2C), being higher during session 2 compared to the others (vs. session 1, p<0.01; vs. session 3, p<0.05). A main effect of stress was also found (F_(1,18)_=13.69, p<0.01), in which STR dams displayed higher AB/LG responses than CTR dams. The session x stress interaction effect was also observed (F_(2,36)_=14.53, p<0.001), with post hoc analysis indicating that STR dams exhibited higher AB/LG responses than CTR dams during session 2 (p<0.001), and that within the STR group, AB/LG was higher during session 2 than during sessions 1 (p<0.001) and 3 (p<0.001).

**Fig2.**
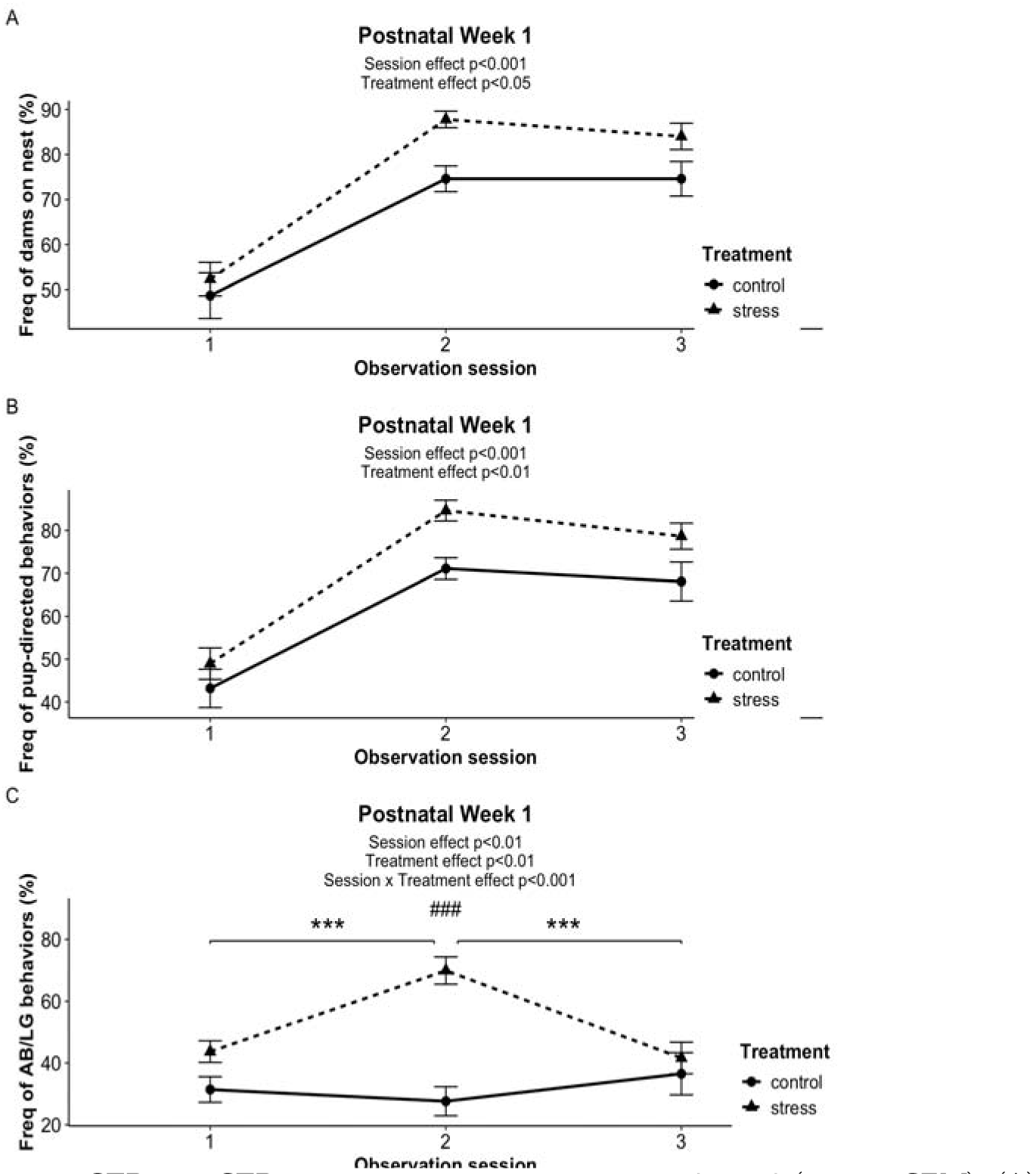
Maternal care outcomes of CTR- and STR-dams in observation sessions 1, 2 and 3 (mean ± SEM). (A) Percentage of frequency of the dam on the nest. (B) Percentage of frequency of pup-directed behaviors. (C) Percentage of frequency of AB/LG. ***p<0.001 indicates vs. session 1 and 3 within the STR group. ###p<0.001 indicates STR vs. CTR in session 2. For the STR group, observation sessions were conducted immediately before (session 1), immediately after (session 2), and 2-3 hr after (session 3) unpredictable maternal separation. CTR: control; STR: stress. (CTR: n=9; STR: n=11).

#### 3.1.4. Cumulative stress effects on SPT

Independent comparisons between CTR and STR dams revealed no significant differences in sucrose preference at any concentration (Suppl. Table S5). Sucrose intake did not differ between groups at 0.5%, 1%, or 2%, while a trend toward higher intake at 4% was observed in STR dams (t_(15.11)_=1.95, p=0.07) (Suppl. Table S5).

### 3.2. Offspring outcomes

#### 3.2.1. Cumulative ELS effect on offspring body weight

Litters were weighed from P1-P14 and the day of weaning (i.e., P21) (Suppl. FigS2). A main effect of day was found (F_(14,252)_=2372.62, p<0.001), with body weight increasing over time in both CTR and STR groups. A day x stress interaction effect was also found (F_(14,252)_=2.08, p<0.05); on average, STR litters were heavier than CTR litters from P1 onward, although *post hoc* tests did not reveal significant differences at individual time points. No main effect of stress was found (F_(1,18)_=1.62, p=0.22). Adult offspring were weighed before each behavioral testing. The 3-way mixed ANOVAs performed by sex showed no main effect of genotype either on males (F_(1,36)_=0.14, p=0.71) or females (F_(1,36)_=0.00, p=0.94); therefore, genotypes were grouped, and 2-way mixed ANOVAs (stress x week) were performed. For males’ body weight (Suppl. FigS3A), a main effect of week was found (F_(6,228)_=288.24, p<0.001), indicating a progressive weight increase. A main effect of stress was also observed (F_(1,38)_=5.48, p<0.05), where STR males weighed more than CTR males. No effect of week x stress interaction was found (F_(6,228)_=0.29, p=0.94). For females’ body weight (Suppl. FigS3B), a main effect of week was found (F_(6,228)_=294.09, p<0.001), with an increase of weight over time. A main effect of stress was also observed (F_(1,38)_=11.12, p<0.01), in which STR females weighed more than CTR females. No week x stress interaction effect was found (F_(6,228)_=0.44, p=0.85).

#### 3.2.2. SERT genotype x cumulative ELS interaction effect on EPM

For distance traveled (Fig3A), no main effect of genotype (F_(1,72)_=0.01, p=0.91), stress (F_(1,72)_=0.01, p=0.89), sex (F_(1,72)_=0.90, p=0.35), or the genotype x stress x sex interaction effect (F_(1,72)_=1.33, p=0.25). No other interaction effects were found either. With regards to the frequency in open arms (Fig3B), a genotype x sex interaction effect (F_(1,72)_=4.12, p<0.05), in which WT-females visited more times the open arms than WT-males (p<0.05), and that HET-females had lower visits compared to WT-females (p<0.05). We also found a 3-way interaction effect of genotype x stress x sex (F_(1,72)_=5.52, p<0.05). Further analysis indicated that HET-STR males visited the open arms more frequently than HET-CTR males (p<0.05); WT-STR females showed a higher frequency compared to WT-STR males (p<0.05); HET-STR females visited the open arms fewer times than WT-STR females (p<0.01); and a trend was found in which HET-STR females had less visits than HET-STR males (p=0.06). No other main or interaction effects were observed. For the time spent in open arms (Fig3C), no main effect of genotype (F_(1,72)_=0.24, p=0.62), stress (F_(1,72)_=0.50, p=0.48), sex (F_(1,72)_=1.20, p=0.28), or the genotype x stress x sex interaction effect (F_(1,72)_=1.32, p=0.25) was found. No other interaction effects were found either.

**Fig3.**
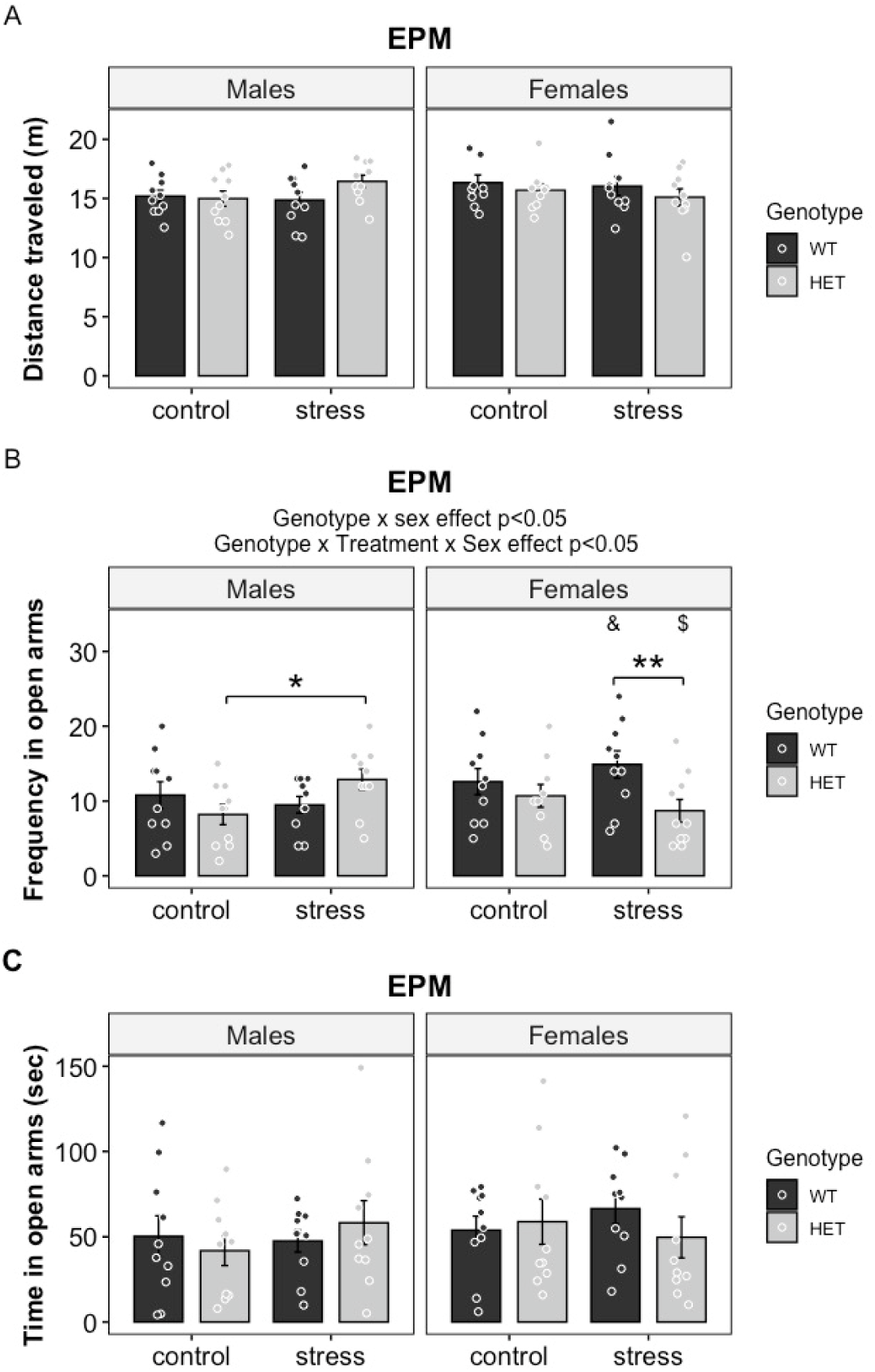
EPM outcomes for WT and HET male and female offspring exposed to control or cumulative ELS conditions. (A) Distance traveled, (B) Frequency in open arms, (C) Time in open arms (mean ± SEM). *p<0.05, **p<0.01; &p<0.05 vs. WT-STR males. $ indicates a trend vs. HET-STR males (p=0.06). EPM: elevated plus maze; HET: serotonin transporter heterozygous knockouts; WT: wildtypes. (n=10 genotype/stress/sex).

#### 3.2.3. SERT genotype x cumulative ELS interaction effects on SPT

Sucrose preference and sucrose intake were analyzed in the SPT, in which animals were exposed to four sucrose concentrations (0.5%, 1%, 2%, 4%). For sucrose preference, separate analyses per concentration revealed no main effects of genotype, stress, sex, or their interactions at any of the sucrose concentrations, except for a trend of sex effect on the preference at 4% (F_(1,72)_=3.33, p=0.07), in which females tended to show a higher sucrose preference than males (Suppl. Table S6). In contrast, the 3-way ANOVA for sucrose intake at 0.5% (F_(1,72)_=22.92, p<0.001), 1% (F_(1,72)_=48.40, p<0.001), 2% (F_(1,72)_=33.60, p<0.001), and 4% (F_(1,72)_=74.63, p<0.001) showed a main effect of sex, indicating that females consumed more sucrose than males regardless of the sucrose concentration (Suppl. Table S7). For the sucrose intake at 2% (Fig4), a marginal effect of stress was found (F_(1,72)_=3.95, p=0.05), in which rats exposed to STR consumed less sucrose than CTR. A genotype x sex interaction effect was also observed (F_(1,72)_=4.15, p<0.05), with further analysis showing that WT and HET females consumed more 2% sucrose than their WT (p<0.05) and HET (p<0.001) male counterparts. No other main or interaction effects were found for any of the other sucrose concentrations.

**Fig4.**
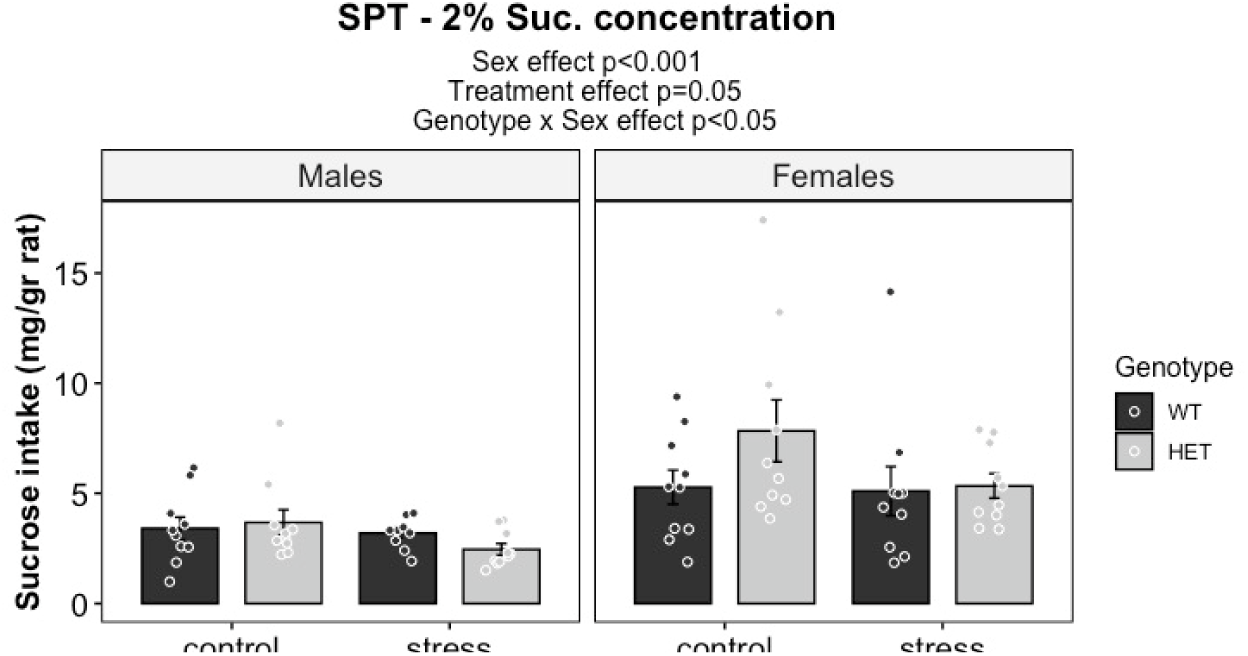
Sucrose intake at 2% concentration of sucrose during the SPT in WT and HET male and female offspring, exposed to control or cumulative ELS conditions. HET: serotonin transporter heterozygous knockouts; WT: wildtypes. (n=10 genotype/stress/sex)

#### 3.2.4. SERT genotype x cumulative ELS interaction effect on OFT

For the distance traveled (Fig5A), a trend of genotype effect was found (F_(1,72)_=3.49, p=0.07), in which HET rats tended to travel longer distances than WT. No main effect of stress (F_(1,72)_=0.47, p=0.49), sex (F_(1,72)_=1.31, p=0.26), or the genotype x stress x sex interaction effect (F_(1,72)_=2.37, p=0.13) was found. No other interaction effects were found either. For the frequency in the center (Fig5B), no main effect of genotype (F_(1,72)_=1.76, p=0.19), stress (F_(1,72)_=0.52, p=0.47), or sex (F_(1,72)_=1.55, p=0.22) was found, but a trend of genotype x sex interaction effect was observed (F_(1,72)_=3.13, p=0.08). The 3-way interaction effect of genotype x stress x sex (F_(1,72)_=4.04, p<0.05) was also observed. HET-STR males visited the center more times than WT-STR males (p<0.01); HET-STR females had a lower frequency compared to HET-STR males (p<0.01); in addition, a trend of HET-STR females visiting the center fewer times in comparison to HET-CTR females (p=0.06), and a trend of WT-STR females visiting the central area more times than WT-STR males (p=0.09) were observed. Time spent in the center (Fig5C) did not differ by genotype (F_(1,72)_=1.34, p=0.25), stress (F_(1,72)_=1.75, p=0.19), or sex (F_(1,72)_=0.70, p=0.41); neither a genotype x stress x sex interaction effect was observed (F_(1,72)_=1.01, p=0.32), nor any other interactions.

**Fig5.**
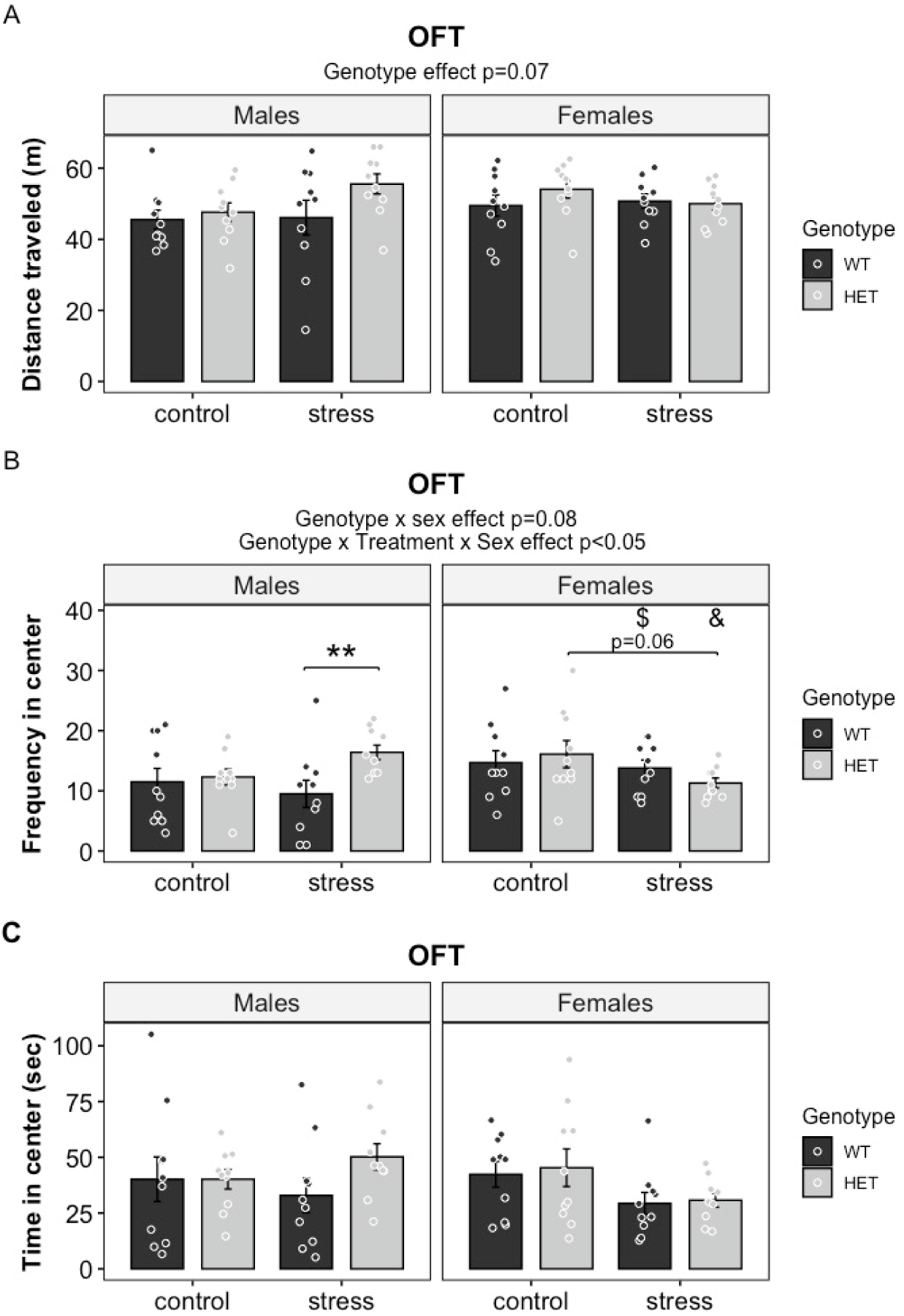
OFT outcomes for WT and HET male and female offspring exposed to control or cumulative ELS conditions (mean ± SEM). (A) Distance traveled, (B) Frequency in center, (C) Time in center. **p<0.01; &p<0.05 vs. STR-HET males. $ indicates a trend vs. STR-WT males (p=0.09). OFT: open field test; HET: serotonin transporter heterozygous knockouts; WT: wildtypes (n=10 genotype/stress/sex)

#### 3.2.5. SERT genotype x cumulative ELS interaction effects on OLT

Ten animals were excluded for not exploring one of the objects during training, and one for no exploration in the memory trial. One-sided t-test comparing each group’s DI to the corresponding chance level (DI_fic)_ indicated a marginally significant effect in which the DI was just above the chance level only in HET-STR females (t_(7)_=1.87, p=0.05). No other one-sided t-test comparisons indicated a DI significantly above the chance level. No main effect of genotype (F_(1,61)_=0.00, p=0.94), stress (F_(1,61)_=0.12, p=0.73), sex (F_(1,61)_=1.35, p=0.25), or the sex x genotype x stress interaction effect (F_(1,61)_=0.55, p=0.46) were observed (Fig6). No other interaction effects were found either.

**Fig6.**
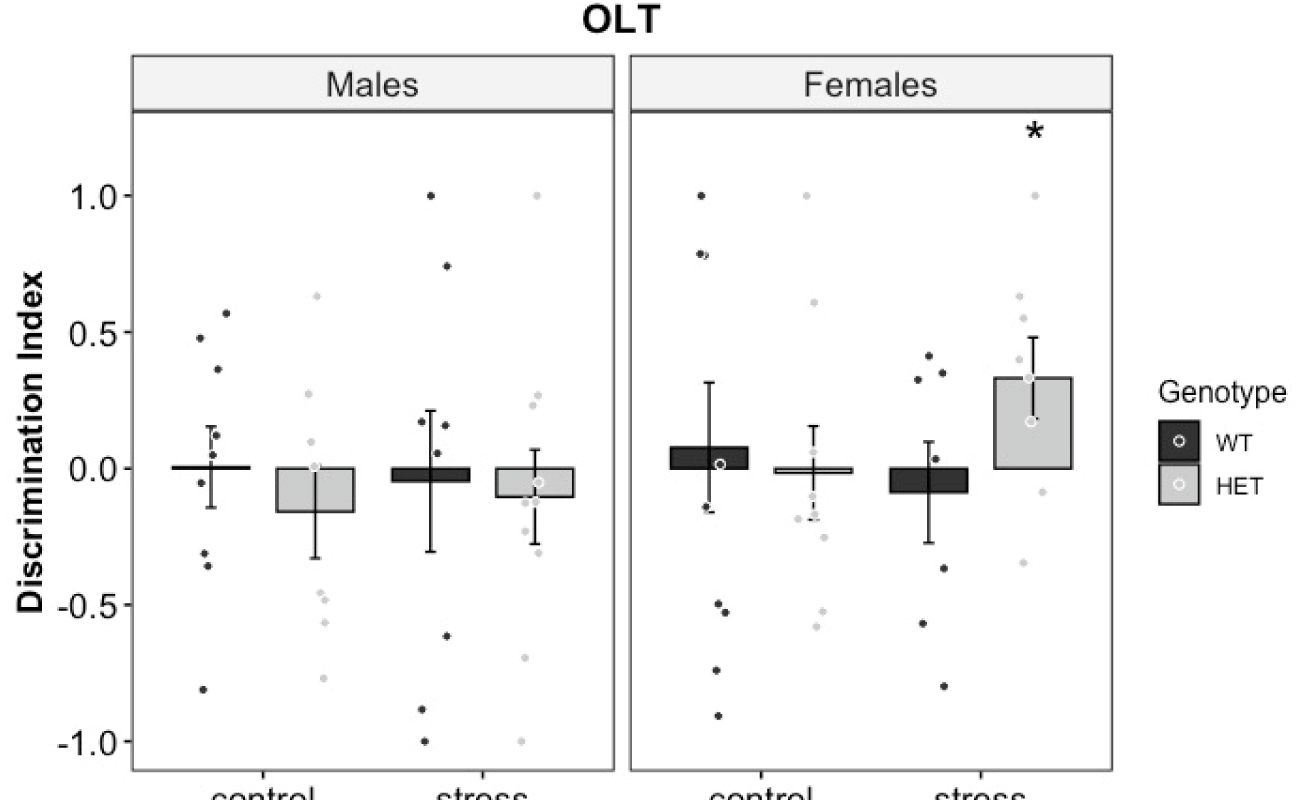
Discrimination index of the novel object location in the OLT for WT and HET male and female offspring exposed to control or cumulative ELS conditions (mean ± SEM). *p=0.05 vs. chance level. HET: serotonin transporter heterozygous knockouts; OLT: object location test; WT: wildtypes (n=7-10 genotype/stress/sex).

#### 3.2.6. SERT genotype x cumulative ELS interaction effect on NOR

Nine animals were excluded for not exploring one of the objects during training, and one for exploring both objects for less than 3 sec, making memory acquisition unlikely. One-sided t-test comparisons between the novel object DI for each group and the corresponding DI_fic_ indicated that the DI was significantly above the chance level in WT-CTR males (t_(9)_=3.87, p<0.01) and WT-CTR females (t_(8)_=2.11, p<0.05); moreover, HET-STR females had a DI significantly above chance level (t_(8)_=2.82, p<0.05), while a trend of higher DI than the chance level was observed in HET-CTR females (t_(8)_=1.70, p=0.06). For none of the other groups, the DI was significantly above the chance level. A main effect of stress on the DI for the novel object was found (F_(1,62)_=4.44, p<0.05; Fig7), where STR rats had a lower DI than CTR rats (p<0.05). We also found the genotype x stress interaction effect (F_(1,62)_=7.07, p<0.01), in which WT-STR rats had a lower DI compared to WT-CTR (p<0.01); a marginal significant effect of higher DI was observed in HET-STR rats compared to WT-STR animals (p=0.05), and a trend of lower DI in HET-CTR animals compared to WT-CTR rats was found (p=0.08). No main effect of genotype (F_(1,62)_=0.03, p=0.87), sex (F_(1,62)_=0.81, p=0.37), or the genotype x stress x sex interaction effect (F_(1,62)_=0.00, p=0.95) was found. No other interaction effects were found either.

**Fig7.**
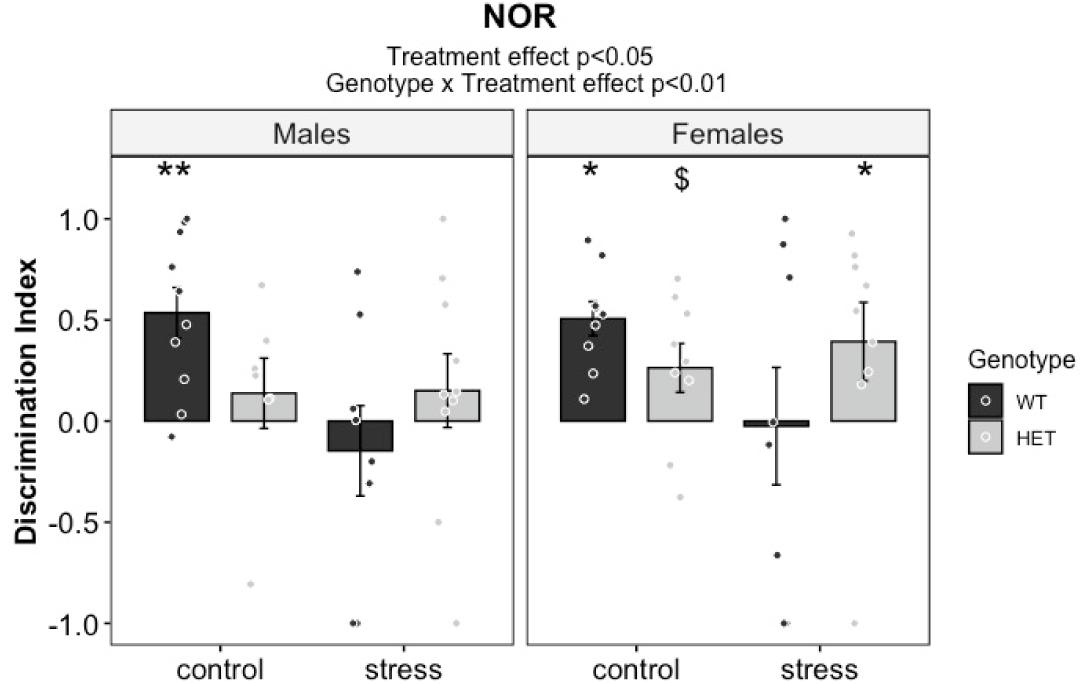
Novel object discrimination index in the NOR for WT and HET male and female offspring exposed to control or cumulative ELS conditions (mean ± SEM). *p<0.05, **p<0.01 vs. chance level. $ indicates a trend vs. chance level (p=0.06). HET: serotonin transporter heterozygous knockouts; NOR: novel object recognition; WT: wildtypes (n=7-10 genotype/stress/sex).

#### 3.2.7. SERT genotype x cumulative ELS interaction effect on SIT

A main effect of sex on social interaction was found (F_(1,32)_=11.59, p<0.01; Fig8), where females exhibited reduced social interaction compared to males (p<0.01). A genotype x sex interaction effect was also observed (F_(1,32)_=5.20, p<0.05), with further analysis revealing that HET-females showed a lower social interaction than HET-males (p<0.001). No main effect of genotype (F_(1,32)_=0.00, p=0.93) or stress (F_(1,32)_=2.46, p=0.13) was found. Neither the genotype x stress x sex interaction effect (F_(1,32)_=0.19, p=0.66) nor the other interactions were significant.

**Fig8.**
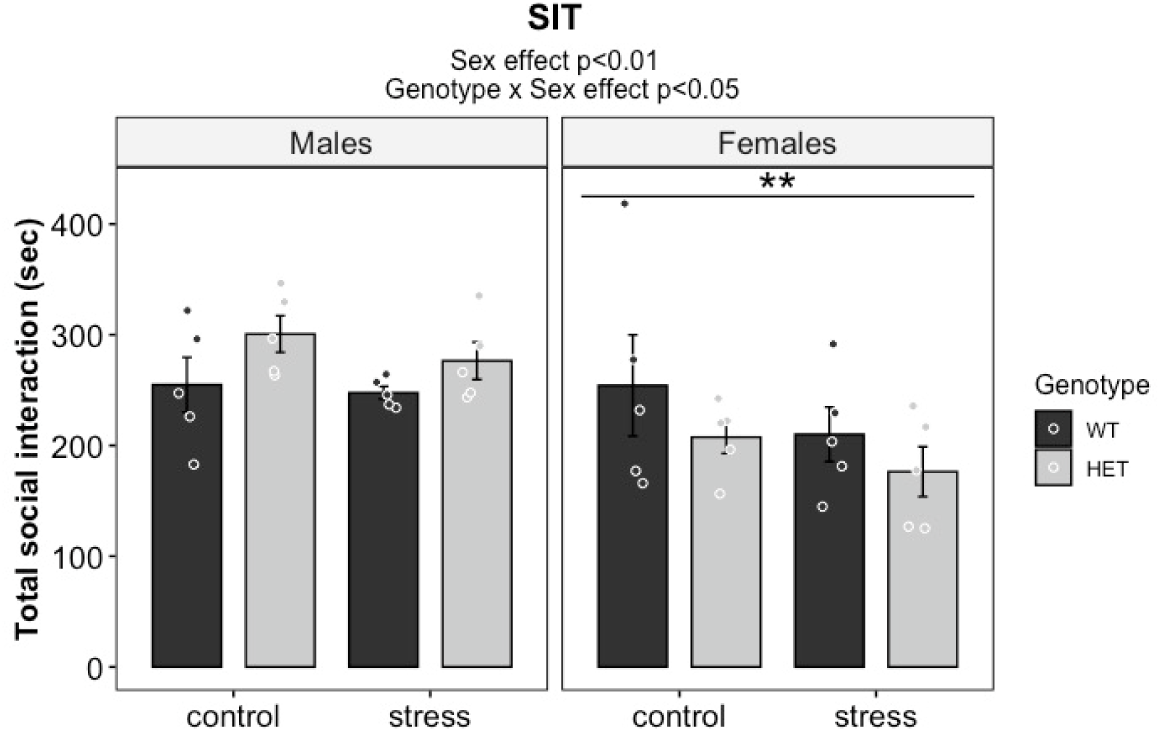
Total time of social interaction in the SIT displayed by WT and HET male and female offspring exposed to control or cumulative ELS conditions (mean ± SEM). **p<0.01 vs. males. HET: serotonin transporter heterozygous knockouts; WT: wildtypes (n=5 pairs genotype/stress/sex).

#### 3.2.8. SERT genotype x cumulative ELS interaction effect on SR

One WT-STR male was mistakenly included twice, and one HET-CTR male was excluded from this analysis as the empty chamber was not visited during the sociability trial. During the sociability trial, the time spent exploring the social stimulus (Fig9A) did not differ significantly by genotype (F_(1,70)_=2.07, p=0.15), stress (F_(1,70)_=0.45, p=0.50), or sex (F_(1,70)_=0.01, p=0.92). Neither a 3-way genotype x stress x sex interaction (F_(1,70)_=0.41, p=0.52) nor other interaction effects were found. No main effects of genotype (F_(1,70)_=0.20, p=0.65), stress (F_(1,70)_=0.06, p=0.80), or sex (F_(1,70)_=0.10, p=0.75), nor any interactions, including genotype x stress x sex (F_(1,70)_=0.29, p=0.60) were found on social preference (Fig9B). During the social recognition trial, the time spent exploring both social stimuli (Fig9C) differed by genotype (F_(1,70)_=8.90, p<0.01), where HET rats explored the social stimuli for longer in comparison to WT (p<0.01). No main effect of stress (F_(1,70)_=0.46, p=0.50) or sex (F_(1,70)_=1.42, p=0.24), or the genotype x stress x sex interaction effect (F_(1,70)_=0.94, p=0.33) was found. None of the other interaction effects were observed either. For the social recognition index (Fig9D), no main or interaction effects were found: genotype (F_(1,70)_=0.28, p=0.60), stress (F_(1,70)_=0.00, p=0.98), sex (F_(1,70)_=0.40, p=0.53), genotype x stress x sex (F_(1,70)_=0.11, p=0.73). No other interaction effects were found either.

**Fig9.**
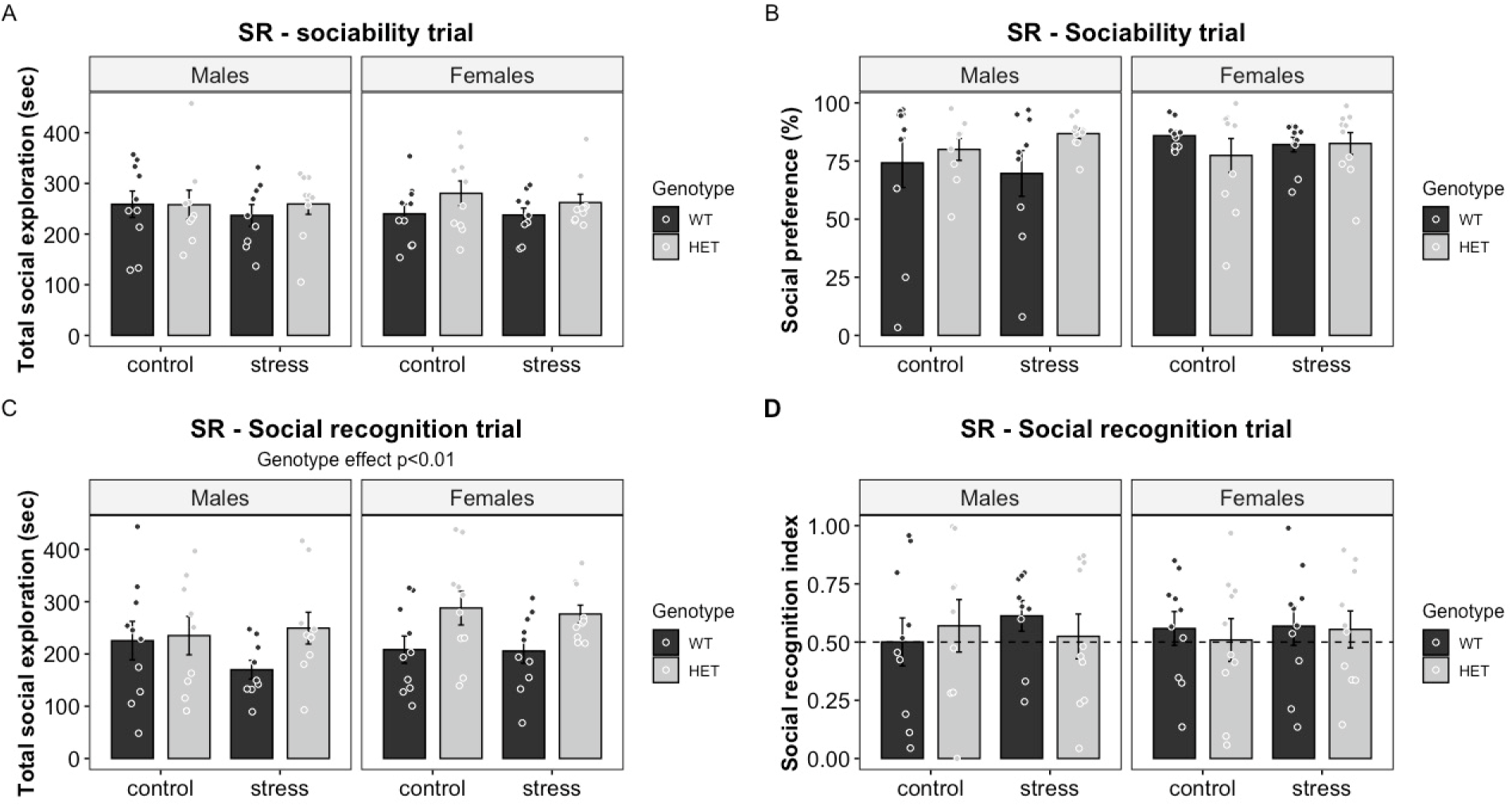
SR outcomes for WT and HET male and female offspring exposed to control or cumulative ELS conditions (mean ± SEM). (A) Total time spent exploring the social stimulus; (B) Social preference; (C) Total time spent exploring both social stimuli; (D) Social recognition index. The long-dashed line indicates no social discrimination. HET: serotonin transporter heterozygous knockouts; WT: wildtypes (n=9-10 sex/genotype/stress).

## 4. Discussion

We aimed to determine whether exposure to *cumulative* ELS induced anxiety-like and depressive-like behaviors in male and female offspring rats. In addition, we examined whether SERT^+/-^ offspring were more vulnerable to cumulative ELS effects than wildtypes. Dams were exposed to stress before, during, and after pregnancy to model the accumulation of ELS for the offspring, and were tested for maternal care and anhedonia. To test potential alterations in affective, cognitive, and social behaviors in the offspring, we assessed them in the EPM, SPT, OFT, OLT, NOR, SIT and SR. Table 1 presents an overview of main and interaction effects. Overall, the accumulation of stress in dams (from the period immediately prior to conception until offspring weaning) enhanced maternal care, increased maternal care towards the pups immediately after UMS, reduced body weight gain, and did not induce anhedonia. The findings from the offspring indicate that: (1) HET animals were more social but also had a tendency toward novelty-induced hyperactivity; HET females exhibited increased anxiety-like behavior, but not HET males; (2) cumulative ELS did not induce a global adverse effect on behavior, but specifically reduced sucrose intake and impaired object recognition; (3) the SERT genotype x cumulative ELS interaction appeared to be beneficial for object recognition; (4) females consumed more sucrose and were less anxious than males, while males exhibited more social interaction than females. In addition, (5) the SERT genotype x sex interaction indicated that the genotype influenced anxiety-like behavior in a sex-dependent manner, increasing anxiety in females but reducing it in males; and (6) the 3-way interaction (SERT genotype x cumulative ELS x sex) revealed that cumulative ELS further amplified the sex-specific effects of genotype, increasing anxiety-like behaviors in HET females, while producing the opposite effect in HET males.

**Table 1.** Overview of genotype, stress, sex and their interaction effects on offspring behavioral outcomes.

|  | **Outcome** | **Main effect of genotype** | **Main effect of stress** | **Main effect of sex** | **Genotype x stress interaction** | **Genotype x sex interaction** | **Stress x sex interaction** | **Genotype x stress x sex interaction** |
| --- | --- | --- | --- | --- | --- | --- | --- | --- |
| **Task** |  |  |  |  |  |  |  |  |
| Elevated Plus Maze (EPM) | Distance traveled | No | No | No | No | No | No | No |
|  | Frequency in open arms | No | No | No | No | **Yes** | No | **Yes** |
|  | Time in open arms | No | No | No | No | No | No | No |
| Sucrose Preference Test (SPT) | Sucrose preference at 0.5% | No | No | No | No | No | No | No |
|  | Sucrose preference at 1% | No | No | No | No | No | No | No |
|  | Sucrose preference at 2% | No | No | No | No | No | No | No |
|  | Sucrose preference at 4% | No | No | **No\*** | No | No | No | No |
|  | Sucrose intake at 0.5% | No | No | **Yes** | No | No | No | No |
|  | Sucrose intake at 1% | No | No | **Yes** | No | No | No | No |
|  | Sucrose intake at 2% | No | **p=0.05** | **Yes** | No | **Yes** | No | No |
|  | Sucrose intake at 4% | No | No | **Yes** | No | No | No | No |
| Open Field Test (OFT) | Distance traveled | **No\*** | No | No | No | No | No | No |
|  | Frequency in center | No | No | No | No | **No\*** | No | **Yes** |
|  | Time in center | No | No | No | No | No | No | No |
| Object Location Test (OLT) | Discrimination index | No | No | No | No | No | No | No |
| Novel Object Recognition (NOR) | Discrimination index | No | **Yes** | No | **Yes** | No | No | No |
| Social Interaction Test (SIT) | Social Interaction | No | No | **Yes** | No | **Yes** | No | No |
| Social Recognition (SR) | Total social exploration (sociability trial) | No | No | No | No | No | No | No |
|  | Social preference (%) | No | No | No | No | No | No | No |
|  | Total social exploration (social recognition trial) | **Yes** | No | No | No | No | No | No |
|  | Social recognition index | No | No | No | No | No | No | No |

\* Indicates a trend

### 4.1. SERT genotype effects

Overall, anxiety-like behavior, anhedonia, object recognition, spatial memory, or social recognition were not consistently affected as main effects of SERT genotype compared to wildtype animals. The literature on anxiety-like, depressive-like, and social behaviors of SERT^+/-^ rodents is mixed. Some studies have shown that SERT^+/-^ animals exhibit increased anxiety-like behaviors (Roversi et al., 2020; Sun et al., 2024) and anhedonia (Houwing, Schuttel, et al., 2020; Sun et al., 2024), while others have observed no changes (Houwing, Schuttel, et al., 2020), even reduced anxiety (Van Den Hove et al., 2011). Other studies did not report anhedonia (Roversi et al., 2020), and even improved stress coping in SERT^+/-^ rodents compared to SERT^+/+^ (Houwing, Schuttel, et al., 2020; van der Doelen et al., 2013). In SERT^+/-^ mice, improved memory performance was reported compared to wildtypes (Van Den Hove et al., 2011). In turn, social interaction has been shown to be reduced in SERT^+/-^ animals (Houwing, Staal, et al., 2019; Roversi et al., 2020). In the present study however, the only main effect of SERT genotype was found on social recognition, where HET rats exhibited increased social investigation when social stimuli were confined (in the SR), but not when freely moving (in the SIT). As we also observed a trend of greater distance traveled in the OFT, we wonder whether HET rats may exhibit higher exploration in novel environments compared to WT, or if they were displaying increased locomotor speed, a sign of hyperactivity (Thompson et al., 2018). We checked *ad hoc* the mean velocity in the OFT, and found that HET rats showed a trend of higher mean velocity in comparison to WT (F_(1,72)_=3.49, p=0.07), thus suggesting a tendency toward increased locomotor activity than exploratory drive per se. In addition, genotype-dependent effects emerged in interaction with sex and/or stress in several tasks (e.g., EPM, OFT, NOR), indicating that SERT-related behavioral differences were context-dependent.. Consistent with this, the evolutionary maintenance of functional 5-HTTLPR variants suggest that each may confer some adaptive advantage in order to persist (Homberg & Lesch, 2011). Yet, the persistence of these variants may also reflect a context-dependent trade-off, such that the s-allele may exert both beneficial and detrimental effects depending on environmental factors. The increased social investigation, together with the trend toward novelty-associated hyperactivity, observed in our SERT^+/-^ rats supports this idea.

### 4.2. Cumulative ELS effects

To our knowledge, this is the first study assessing the behavioral phenotype of rats exposed to cumulative ELS to mimic the mother-to-offspring continuity of adversity exposure. Unexpectedly, the combined exposure to maternal pregestational and prenatal stress with postnatal ELS did not produce a robust maladaptive behavioral phenotype in the offspring; instead, it produced selective effects, including reduced sucrose intake and impaired object recognition.

STR animals consumed less 2% sucrose solution than CTR.Although this may reflect a diminished drive to consume sugar, the sucrose preference did not change, suggesting that a hedonic responding (“liking”) remained intact. One alternative is that metabolic changes induced by cumulative ELS played a role in the regulation of sucrose intake, given the known influence of stress exposure on energy balance (Ulrich-Lai & Ryan, 2014). Even though the difference between STR and CTR rats was only statistically significant at 2%, STR rats showed lower sucrose intake at the other concentrations as well (data not shown). Similar to other studies exposing animals to prenatal (Abramova et al., 2021) and postnatal ELS (Alcántara-Alonso et al., 2017; Colleluori et al., 2022; Foright et al., 2024), STR offspring were persistently heavier than CTR at each measured time point (Suppl. FigS2), with the weight difference reaching statistical significance at adulthood (Suppl. FigS3). Thus, cumulative ELS may have increased the STR rats’ metabolic risk, potentially causing alterations in sucrose metabolism that influence their sugar intake, similar to what has been reported in animals exposed to postnatal ELS (Ruiz et al., 2018; Turner et al., 2024).

Cumulative ELS also impaired object recognition in STR rats compared to CTR, an effect reported before in rodents exposed to prenatal (Cabrera et al., 1999; Fidilio et al., 2022; Ivanova et al., 2023) and postnatal ELS (Kotlinska et al., 2023; Reincke & Hanganu-Opatz, 2017; Talani et al., 2023). Performance in the NOR has been associated with hippocampal function (Furini et al., 2020; Tuscher et al., 2018), which is known to be particularly vulnerable to prenatal (Buck et al., 2024; Soti et al., 2022) and postnatal ELS (Brosens et al., 2023; Dayananda et al., 2023; Hofstra et al., 2024). Another study linked impaired object recognition to prenatal ELS, mediated by reduced cAMP response element-binding protein in the hippocampus (Ivanova et al., 2023), while impaired object recognition in rats exposed to postnatal ELS was associated with reduced hippocampal volume and altered levels of GABA, glutamate, glutamine, and glycine in this region (Kotlinska et al., 2023). Therefore, hippocampal dysfunction may underlie the impaired object recognition of animals exposed to cumulative ELS, although the specific mechanism involved remains to be demonstrated. Using the OLT, we expected to further confirm altered spatial memory in STR rats compared to CTR, an ability also dependent on the hippocampus. However, rats without any intervention (WT-CTR males and females) did not show a DI significantly above chance level, suggesting that the task did not effectively assess spatial memory at baseline conditions. This could be due to poor visibility of spatial cues under dim lighting which may have increased the task difficulty, or because the training trial was too short for the animals to learn the objects’ positions, although a 3-min training trial has previously been shown to be sufficient for acquisition of object memory (Olivier et al., 2008, 2009).

Remarkably, STR-dams stayed on the nest and showed pup-directed behaviors with more frequency than CTR-dams, and intensified their active maternal care immediately after pup reunion following UMS. Moreover, STR-dams did not exhibit anhedonia. We speculate that females exposed to stress immediately prior to conception may have developed a form of stress adaptation during pregnancy and lactation, potentially due to familiarity with certain stressors. From G14-G20, dams were restrained in transparent cylinders, the same apparatus used for restriction during the pregestational stress protocol. Postnatally, within the UMS timeframe, dams were either placed in the same cylinders or exposed to forced swimming; again, stressors they had previously encountered. Although unpredictability was introduced as an additional source of stress by randomizing the daily order of stressors for each dam and the timing of each stressor, prior familiarity with the stressors themselves may still have contributed to dams’ adaptation to chronic stress (Parihar et al., 2011). It is also possible that the higher frequency of active maternal care in STR-dams, including AB/LG nursing and pup-directed behaviors such as nest building, reflects a proactive coping style. This interpretation is consistent with evidence that rodents exhibiting proactive stress-coping strategies often display nest-building and home-territory defense (Koolhaas et al., 1999). In addition, STR-dams lost weight during pregestational and prenatal stress, but this effect was not sustained into the postnatal stress period. This finding may further support adaptation of STR-dams to chronic stress, such that they were resilient when tested in the SPT, thus explaining the lack of differences with CTR-dams. Overall, the accumulation of stress in dams was associated with enhanced aspects of maternal care, did not appear to impair with reproduction, and did not induce anhedonia.

It is worth noting that no anhedonia, increased anxiety, or impaired social behaviors were observed in offspring exposed to cumulative ELS, despite the fact that prenatal and postnatal ELS commonly induce altered affective and social behaviors in the offspring (for reviews see: Lupien et al., 2009; Weinstock, 2017). The outcomes limited to sucrose intake and object recognition suggest that pregestational stress exposure did not amplify the anticipated negative effects of prenatal and postnatal ELS. A previous study showed that the combination of prenatal and postnatal ELS did not increase anxiety-like behaviors, impair the social recognition, or alter spatial memory (Corredor et al., 2022), while pregestational stress was found to reduce anxiety-like responses and did not induce depressive-like behavior in rats (Houwing, Schuttel, et al., 2020). Mechanistically, one speculation is that cumulative ELS did affect the developing brain but that the increased maternal care shown by STR-dams buffered some of these effects.

### 4.3. SERT genotype x cumulative ELS effects

The original Caspi’s study of the 5-HTTLPR x stress interaction highlighted that gene-environment interactions are critical for understanding vulnerability to human depression, with accumulation of stressful experiences playing an important role (Caspi et al., 2003). In our study, we modeled the accumulation of stressors in early life by designing a cumulative ELS paradigm that integrated maternal stress as an earlier stressor, and compared its effects between wildtypes and heterozygous SERT knockouts. This model did not produce wide-spread alterations in affective-related behaviors. Instead, we found that HET-STR animals outperformed their WT-STR counterparts in the NOR, while no other consistent genotype x stress effects were observed across behavioral tasks. Thus, it appears that the SERT^+/-^ genotype may confer a selective advantage for object recognition under cumulative ELS, without clear detrimental effects on other behavioral domains.

Cognitive outcomes in SERT^+/-^ rodents exposed to ELS have rarely been examined. Existing studies report no detrimental effect of the SERT genotype x prenatal ELS interaction on object memory (Van Den Hove et al., 2011), and intact social recognition in SERT^+/-^ females exposed to postnatal ELS (Houwing, Ramsteijn, et al., 2019). It is unclear whether and how the lifelong reduction in SERT expression impacts hippocampal functioning and thereby cognitive performance in SERT^+/-^ rodents. While hippocampal SERT expression of SERT^+/-^ rats is reduced by 40-50% (Homberg et al., 2007), serotonin uptake in brain slices and serotonin extracellular levels remain unchanged (Homberg et al., 2007; Olivier et al., 2008). Therefore, it remains to elucidate which molecular pathways mediate the interaction between reduced SERT expression and cumulative ELS in shaping hippocampal-dependent functions, such as those assessed in the NOR. Interestingly, a previous study explored the SERT genotype x ELS interaction on the transcription of glucocorticoid receptors (i.e., GR and MR) in the hippocampus as a potential mechanism underlying the onset of depressive symptoms (Van Der Doelen et al., 2014). This investigation was based on the hypothesis that affective and cognitive alterations in depression may be linked to altered function of the ventral and dorsal hippocampus, respectively (Fanselow & Dong, 2010). van der Doelen et al., 2014 found that SERT^+/-^ rats exposed to postnatal ELS had increased mRNA expression of MR in both the dorsal and ventral hippocampus. However, in our study, where animals were exposed to cumulative ELS, it remains unclear whether similar mechanisms are involved, or whether maternal care also played a role, since STR-dams showed higher nest frequency, more pup-directed behaviors, and increased AB/LG. As offspring of high AB/LG dams exhibit enhanced memory linked to an increased hippocampal GR expression (Liu et al., 2000), it is relevant to consider whether the improved memory performance of SERT^+/-^ rats exposed to cumulative ELS was mediated by differential MR and/or GR hippocampal activity induced by cumulative ELS, high maternal care, or both.

Of note, a literature review on ELS effects in SERT^+/-^ rodents concluded that low maternal care is one factor that appears to increase susceptible to anxiety- and depressive-like behaviors (Houwing et al., 2017). Indeed, it was previously shown that SERT^+/-^ animals receiving low maternal care, but not wildtypes, exhibited enhanced anxiety-like responses (Carola et al., 2008). In turn, high maternal care increased serotonin levels and decreased serotonin turnover in the hippocampus of SERT^+/-^ offspring, bringing them to levels similar to those observed in wildtypes (Carola et al., 2011), which may indicate a potential buffering or rescue effect of high-quality maternal care in SERT^+/-^ offspring. Therefore, we propose that the higher quality of maternal care provided by STR-dams may have been particularly beneficial for the SERT^+/-^ offspring exposed to cumulative ELS, potentially contributing to improved memory performance while protecting against the emergence of increased anxiety, anhedonia, or social impairments. However, this interpretation remains speculative and requires direct mechanistic investigation. It remains to be determined whether the mechanisms involved are related to an increased hippocampal GR expression, modulation of serotonergic signaling, their interaction, or other processes. For instance, increased GR expression in SERT^+/-^ offspring of well-caring STR-dams might activate molecular pathways such as brain-derived neurotrophic factor, which is crucial for regulating the serotonin system (Homberg et al., 2014). Through such pathways, changes in hippocampal activity might mediate enhanced stress adaptation or cognitive performance in SERT^+/-^ offspring exposed to cumulative ELS and receiving high-quality maternal care.

### 4.4. Main sex-specific effects

Incorporating behavioral analysis by sex is fundamental for understanding the biology of mental health and disease, and for identifying early, sex-dependent risk factors (Beery & Zucker, 2011). Consistent with previous reports, we also found that female rats consumed more sucrose than males (Houwing, Schutter et al., 2020; Grim et al., 2022), while males engaged in more social interaction than females (Karlsson et al., 2015; Peña et al., 2006). From an evolutionary perspective, male and female behavioral adaptations differ in domains where they face different challenges for survival and reproduction (Palanza, 2001). In support of this idea, greater sugar intake in females may reflect an adaptive mechanism to support the high energy demands of reproduction and lactation, and to ensure adequate fat storage for pregnancy. This may be linked to sex differences in glucose homeostasis, which may also place females at a higher risk for obesity and type 2 diabetes (Mauvais-Jarvis, 2024). A meta-analysis of human studies found that total dietary consumption of sugar was associated with an increased risk of depression in the general population (Xiong et al., 2024). Although extrapolation from rodent to humans should be made with caution, this could suggest that higher sugar consumption in females may be relevant in the context of their increased vulnerability to depression, consistent with the higher prevalence of depression observed in women (Albert, 2015). With respect to enhanced social interaction in males, our results are consistent with findings showing that males exhibit higher social engagement towards same-sex conspecifics than females (Douglas et al., 2004; Johnston & File, 1991). Presumably, social stimuli are more rewarding for males than for females, as suggested by increased dopamine activity in the amygdala of males in response to same-sex peer interaction, an effect not observed in females (Weiss et al., 2015). In line with this, after one week of social isolation, males showed increased social reward-seeking behaviors, whereas females exhibited increased sucrose reward-seeking behaviors (Isaac et al., 2024).

As others have reported, we also found no sex differences in sucrose preference (Bruijnzeel et al., 2019; Fonseca-Rodrigues et al., 2022), object recognition or object location (see for a review: Becegato and Silva, 2022), and social preference or social cognition (Bluthé & Dantzer, 1990; Karlsson et al., 2015; Netser et al., 2017; Veenema et al., 2012). Although no main effect of sex emerged on anxiety-like responses, the *post hoc* analysis of the genotype-by-sex interaction did reveal that WT females had more entries into the open arms of the EPM and central area of the OFT than WT males. This suggests that females, at least in the absence of SERT disruption, display reduced anxiety-like responses compared to males, as previously reported (Börchers et al., 2022; for a review: Lovick & Zangrossi, 2021). Given the idea that greater female variability precluding the study of sex differences has been disproven (Kaluve et al., 2022), alternative explanations must be considered to advance our understanding of sex as a biological factor. One possibility is that lower baseline anxiety in females reflects a typical sex difference in rodent models (Lovick & Zangrossi, 2021). Alternatively, standard behavioral paradigms, originally validated in males, may incompletely capture female-typical anxiety responses (Lovick & Zangrossi, 2021).

### 4.5. SERT genotype x sex effects

Several of our findings indicate that reduced SERT expression primarily modulates anxiety in a sex-dependent manner. On one hand, HET females made fewer entries into the EPM open arms than their WT counterparts, a pattern that was absent in HET males, who made a similar number of open-arm entries as their WT counterparts. In addition, HET males entered the open field center more often than WT males, an effect not found in females. On the other hand, HET females showed lower social interaction than HET males in the SIT. This last finding can be interpreted as increased social anxiety, as the time interacting with a partner is influenced by anxiety-related states (Lezak et al., 2017). It appears that lifelong low SERT expression is associated with exacerbated anxiety-like behaviors in females but not in males, which is consistent with previous findings showing that SERT^+/-^ females, but not SERT^+/-^ males, exhibit higher levels of anxiety and depressive-like responses (Houwing, Schuttel, et al., 2020; Van Den Hove et al., 2011). Possibly, these findings are linked to differences in activity of emotional brain circuits between males and females with reduced SERT expression. Human and rodent studies consistently show that the amygdala plays a central role in emotion regulation (Andrewes & Jenkins, 2019). As human carriers of the s-allele exhibit increased activation of amygdala during emotional processing (Kobiella et al., 2011), it has been proposed that reduced SERT expression may contribute to increased amygdala activity, thereby contributing to s-allele carriers’ susceptibility to affective disorders (Loewenstern et al., 2019; Schipper et al., 2019). Considering that brain serotonin levels and SERT sensitivity differ by sex (Carlsson et al., 1985; Nishizawa et al., 1997), it is possible that sex-dependent mechanisms influence how lifelong reduced SERT expression affects amygdala function, potentially contributing to heightening anxiety levels in females but not in males. Indeed, a study reported sex-specific amygdala reactivity influenced by the SERT genotype in mice, although this effect was observed in males but not in females, where increased amygdala reactivity to predator odor was modulated by the SERT genotype (Kolter et al., 2021). This discrepancy suggests that the direction of sex-specific effects may depend on the context or behavioral domain assessed.

The SERT genotype x sex interaction effect on 2% sucrose intake revealed that both WT and HET females consumed more 2% sucrose solution than their respective male counterparts, suggesting the SERT genotype effect as the main driver. We wondered whether the magnitude of these differences was greater between HET males and females than between WT males and females, so that we examined *ad hoc* both effect sizes. Hedges’ *g* results showed a larger difference between HET males and females (Hedges’ *g* = 1.29, 95% CI = [0.98, 1.60]) compared to WT males and females (Hedges’ *g* = 0.59, 95% CI = [0.34, 0.84]), suggesting that reduced SERT expression may amplify sex differences in sucrose intake. In line with the idea mentioned above that higher sugar consumption in women might be linked to increased vulnerability to depression, one might speculate that the SERT genotype modulates sex-specific regulation of reward-related consumption. However, this interpretation remains speculative, and more studies directly comparing SERT^+/-^ males and females are needed to clarify the extent to which reduced SERT expression moderates sex-dependent risks for depression.

The effect of SERT genotype on performance did not differ between males and females in the OLT, NOR, and SR tasks. Studies assessing the behavior of both SERT^+/-^ males and females are limited, and existing research has focused primarily on affective behaviors in tasks like the EPM, OFT, SPT (Joeyen-Waldorf et al., 2009; Houwing, Schuttel et al., 2020; Houwing, Staal et al., 2019; Willadsen et al., 2021). One study reported no SERT genotype x sex interaction on cognitive performance in the NOR (Van Den Hove et al., 2011); another found reduced social preference in SERT^+/-^ males and females, although they were compared to wildtypes and not to each other (Tanaka et al., 2018). Additionally, some studies combined SERT^+/-^ males and females into one group to compare outcomes from object recognition and social interaction with wildtypes, implying no differences between SERT^+/-^ males and females (Kalueff et al., 2007).

Overall, reduced SERT expression appears to modulate affective-related behaviors in a sex-specific manner, while having no observable impact on cognitive performance or social cognition in SERT^+/-^ males and females. These findings underscore the importance of considering gene-by-sex interactions when investigating the underpinnings of anxiety and depression, and highlight the need for more studies examining whether sex-specific vulnerability to affective disorders is linked to such interactions involving the SERT genotype.

### 4.6. The triple SERT genotype x cumulative ELS x sex interaction

Some of the triple SERT genotype x cumulative ELS x sex interaction findings in the EPM and OFT suggest that cumulative ELS may differentially modulate anxiety-like behavior depending on sex in SERT^+/-^ rats, increasing it in females while reducing it in males. HET-STR females were more anxious than HET-CTR females (a trend was observed in the OFT) and compared to WT-STR females (in the EPM). In addition, HET-STR females were more anxious than HET-STR males in both tests. In turn, HET-STR males were less anxious than HET-CTR males (in the EPM) and than WT-STR males (in the OFT). As these effects were not consistently observed across all anxiety-related measures, they should be interpreted with caution. A sex-specific effect of the SERT genotype x ELS interaction has been reported in a limited number of studies assessing affective-related response, with mixed findings. SERT^+/-^ female offspring exposed to prenatal ELS tested showed higher anxiety when tested in the elevated zero maze, an effect not observed in prenatally stressed SERT^+/-^ males (Van Den Hove et al., 2011). The opposite pattern was found in another study where SERT^+/-^ female offspring exposed to prenatal ELS exhibited lower anxiety compared to prenatally stressed wildtype females, an outcome that was not observed in males (Woo et al., 2023). In another study, pregestational stress induced anhedonia in SERT^+/-^ females but not in SERT^+/-^ males; however, no increase in anxiety was observed in either sex (Houwing, Schuttel, et al., 2020).

As discussed above, elevated anxiety in females with reduced SERT expression may be linked to an altered neural circuitry involved in emotion regulation. It is therefore possible that cumulative ELS further modulates these circuits, thereby contributing to higher anxious-like responses in SERT^+/-^ females exposed to cumulative ELS compared to both wildtypes and SERT^+/-^ males. Both prenatal and postnatal ELS induce lasting changes in the hypothalamic-pituitary-adrenal (HPA) axis, including modifications in the expression of HPA-related hormone receptors (e.g, MR, GR, CRHR) in the amygdala, prefrontal cortex, and hippocampus (Van Bodegom et al., 2017). Although we cannot determine whether cumulative ELS adds to the effects of prenatal and/or postnatal ELS, it cannot be ruled out that cumulative ELS may alter the expression of such HPA-related hormone receptors.

We proposed in section 4.3 that high-quality maternal care of STR-dams could have protected SERT^+/-^ rats exposed to cumulative ELS from increased anxiety levels, anhedonia, and social impairments that might have emerged otherwise. Including sex in the SERT genotype x cumulative ELS analysis raises the question of whether SERT^+/-^ females benefited less from high-quality maternal care, while SERT^+/-^ males may have benefited more. Some evidence showing sex-specific effects on maternal care on adult offspring behavior might point in this direction. Increased maternal care in dams following pup reunion after maternal separation was associated with poorer memory function and altered transcriptomics in the medial prefrontal cortex of female, but not male mice (Orso et al., 2025). Conversely, reduced maternal care altered spatial memory and reversal learning only in males (Mooney-Leber & Brummelte, 2020). Moreover, fragmented maternal care induced by limited bedding and nesting led to lasting impairments in spatial abilities across developmental stages in males, whereas in females, these effects were more transient (Bath et al., 2017). However, as these studies were conducted in wildtype animals, their relevance to SERT^+/-^ models remain uncertain, and further research is warranted to elucidate the specific effects of maternal care on SERT^+/-^ offspring and to determine whether these effects differ by sex.

### 4.7. Limitations

It is important to acknowledge some limitations within our study. First, although our cumulative ELS paradigm was designed to mimic the intergenerational continuity of psychosocial adversity, its translational relevance to the human condition remains limited. Despite the introduction of unpredictable pregestational, prenatal, and postnatal stressors to better approximate the uncontrollable nature of real-life adversity, the stressors applied (e.g., food restriction, restraint, forced swim) still represent discrete, experimenter-controlled physical challenges that differ from the psychosocial stressors typically experienced by humans and are inherently difficult to reproduce in animal models. Second, the partial overlap of stressor types across stress exposure periods may have induced habituation or stress adaptation in dams, rather than chronic adversity, thereby potentially limiting the emergence of adverse behavioral outcomes. The observation that STR-dams increased pup-directed behaviors suggests a potential adaptive response to repeated stress. Third, without assessments during earlier developmental stages (juvenile or adolescent), it remains unclear whether behavioral alterations emerged gradually or reflect long-term adaptations. This limits conclusions about when SERT genotype, ELS-, or sex-related changes manifest and whether compensatory mechanisms develop across ontogeny that contribute to the outcomes observed in adulthood. Fourth, the absence of intermediate experimental groups isolating specific stress periods (e.g., prenatal + postnatal, or pregestational + prenatal exposure) constrains interpretation of whether the observed outcomes truly reflect cumulative effects or are driven by a particular developmental window of vulnerability. Including such groups would clarify potential non-linear interactions between stress exposures and identify critical periods to study temporal specificity in stress-related programming. Fifth, the sample size per subgroup (n≈10 per genotype × stress × sex) may have limited statistical power to detect small or interaction effects, particularly given the complexity of three-way designs. This may have contributed to the presence of trends rather than statistically significant findings in several outcomes. Lastly, the OLT failed to elicit discrimination performance above chance even in control animals, indicating limited task sensitivity to detect spatial memory effects under the current conditions, consistent with a finding previously reported by our group (Houwing et al., manuscript in preparation). Future work should optimize OLT parameters, such as longer training, enhanced salience or distal spatial cues, and improve lighting conditions, to more reliably assess spatial memory function in rats.

### 4.8. Conclusion

Our study demonstrates that reduced SERT expression did not consistently increase vulnerability to cumulative ELS across affective, cognitive and social behaviors. Rather, its effects appear to be subtle and context-dependent, including increased social investigation and greater susceptibility to anxiety-like behavior, especially in SERT^+/-^ females. The cumulative ELS paradigm, spanning stress exposure from pregestational to postnatal period to simulate a mother-to-offspring continuity of adversity, did not induce widespread affective-related impairments in the offspring, but instead produced selective effects on sucrose intake and object recognition. Remarkably, accumulation of stress in dams was associated with increased maternal care, promoted active maternal care following UMS, and did not induce anhedonia, suggesting adaptive responses to repeated stress exposure. The SERT genotype x cumulative ELS interaction was associated with improved object recognition performance, while anxiety-like responses were modulated in a sex-dependent manner. SERT^+/-^ females tended to exhibit increased anxiety-like behavior, whereas SERT^+/-^ males showed the opposite pattern in specific contexts. In advancing the use of rodent models to study the biological underpinnings of the 5-HTTLPR x stress interaction associated with a higher risk for affective disorders, our findings highlight the importance of considering sex and environmental context as critical modulators of genotype effects.

## Funding

This work was supported by the Colombian Department of Science, Technology and Innovation (MINCIENCIAS, scholarship program No. 860) but it had no role in study design, data collection, data analysis, data interpretation, or the writing of the report.

## CRediT authorship contribution statement

**Mayerli A. Prado-Rivera:** Conceptualization, Funding Acquisition, Data Curation, Investigation, Formal Analysis, Writing – original draft. **Annemarijn MJ. Fortuin:** Investigation, Data Curation. **Cato M. Drion:** Writing – review & editing**. Jocelien D.A. Olivier:** Conceptualization, Supervision, Writing – review & editing. All authors approved the final version of the manuscript.

## Declaration of Generative AI and AI-assisted technologies in the writing process

During the preparation of this work, MAPR used Alphabet’s Gemini in order to assist with readability and language editing. After using this tool, MAPR reviewed and edited the content as needed and takes full responsibility for the content of the publication.

## Declaration of Competing Interest

The authors have no conflict of interest to declare.

## Data availability

Data will be made available on request.

## Suppl. File 1

### Methods

### Pregestational stress

STR-females were individually housed and subjected to three weeks of CUS, consisting of exposure to 1-2 stressors per day, similar to Gemmel et al., 2019 (Suppl. Table S1)

### Prenatal stress

STR-dams were subjected to unpredictable repeated episodes of restraint stress in a transparent cylinder (20 cm length□×□9□cm diameter□×□9□cm height) adjustable for animal’s size, under a bright light for 45 min three times daily, from G14 to G20. URS was administered during the dark part of the reverse light/dark cycle, keeping a timeframe of 2 hr minimum between restraint episodes. Dams were daily weighed to assess stress effect on weight gain during pregnancy. Dams were left undisturbed from G21 until delivery.

### Breeding

The day after the end of the last stressor, estrous cycle was measured in all females with an impedance apparatus (model MK-11, Muromachi, Tokyo, Japan). When the apparatus indicated the proestrus stage (≥ 3 kiloohms (kΩ)), 1-2 receptive females were housed overnight with one Wistar SERT+/- (Slc6a41Hubr) male from our colony to produce SERT+/+ wildtype (WT) and SERT+/- (HET) male and female offspring. The next day was considered gestational day (G)1 and each female was individually housed, and nest material was provided. Females were left undisturbed until prenatal ELS was applied during the last week of pregnancy.

### Postnatal stress

Dams were checked daily at 9:00 AM and 5:00 PM for delivery, which was set as postnatal day (P)0. From P1-P14, STR-dams and their offspring were daily exposed to unpredictable stress. Pups were exposed to 3 hr of unpredictable maternal separation (UMS), applied at unpredictable time points each day, during the dark part of the reverse light/dark cycle. During UMS, the whole litter was removed from the home cage and placed in preheated Makrolon type 2 cages (P1-P7: 32 ± 1□C; P8-P14: 28 ± 1□C). Somewhere within the 3hr of UMS timeframe, a single stressor was randomly applied to the dam (20 min of restraint stress or 5 min of forced swimming). For restraint stress, the same cylinder as for prenatal ELS was used. For the forced swimming, the dam was placed in a cylindrical Plexiglas tank (50×18 cm diameter) filled up with 30 cm of water at 18 °C ± 2□C. CTR pups were handled for 15 min to control for handling effects. Around P14, pups were ear clipped for genotyping (genotyping procedure: El Aidy et al., 2017). Dams and litters were daily weighed between P1-P14 and in the weaning day (i.e., P21). As adults, offspring were weighed before each behavioral testing.

#### Behavioral testing in adult offspring

### Elevated Plus Maze (EPM)

The EPM consisted of 2 open (45×10×1 cm) and 2 closed (45×10×50 cm) arms opposite to each other elevated at 50 cm. The rat was placed in the center of the maze facing an open arm and was allowed to freely explore the maze for 5 min. Behavior was recorded using automated animal tracking software (EthoVision XT11, Noldus, The Netherlands) to calculate the total distance moved (m) and the frequency and time spent on the open arms.

### Sucrose Preference Test (SPT)

Animals were individually housed and habituated to two bottles of water for 48 hr. The same procedure of SPT used for dams, presented in main text (section 2.3.2) was followed to test the offspring. Sucrose preference and sucrose intake were calculated using the formulas to calculate the same outcomes in dams. After testing, rats were socially housed for the rest of the study.

### Open Field Test (OFT)

The OFT consisted of an open square arena (100×100×40 cm), with a black wooden floor and walls. The arena was subdivided into a center (1 square of 60×60cm) and outer area near the walls (20 cm wide). Behavior was recorded for 10 min using automated animal tracking software (EthoVision XT11, Noldus, The Netherlands) to calculate the total distance moved (m) across the arena, as well as the duration and frequency in the center area.

### Object Location Test (OLT)

The OFT was deemed as habituation for this task. The day of testing, before the training trial the rat was allowed to explore the open arena for 3 min as rehabituation. Then, the animal was removed to a waiting cage (Makrolon type 3 cage: 38.2×22.0×15.0 cm), the arena cleaned, and two identical objects (smooth glass, sand clock-shape bottles) were placed in two opposite corners. The rat was placed back into the open arena and allowed to explore the objects for 3 min, after which the animal was placed in the waiting cage, the arena and objects cleaned, and a second set of identical objects was placed for the memory trial. In this trial, one of the objects was relocated to another corner. The animal was allowed to explore the objects for another 3 min, after which the rat was returned to its home cage. Object’s exploration for both trials was automatically tracked by using EthoVision XT18 software (Noldus, The Netherlands). Object exploration was defined as the rat’s nose pointing towards the object from a distance of 2 cm or less, or directly touching the object. Exploration time of identical objects (obj1, obj2) in the training trial, and exploration time of objects in the familiar (fam loc) and novel (new loc) location in the memory trial were scored. Total time of exploration during memory trial was calculated as: new loc + fam loc, and the discrimination index (DI) of the novel object location was calculated as:

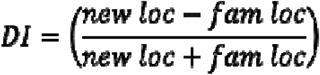

A DI above zero (i.e., chance level) indicated discrimination of the object in the novel location.

### Novel Object Recognition (NOR)

The same general procedure for the OLT was followed to test the novel object recognition and track objects exploration. For this task, the training trial consisted of exploration of two identical objects (wrinkled, glass bottles) placed in two opposite corners of the open arena. The memory trial consisted of exploration of one familiar and one novel object (smooth glass, multiple-marble-shape bottle) placed in the same position as in the training trial. For the memory trial a second set of objects was used. Based on the time exploring the familiar (F) and the novel (N) objects during the memory trial, the DI of the novel object was calculated as:

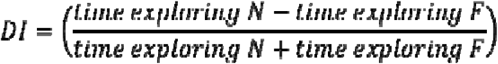

A DI above zero (i.e., chance level) indicated discrimination of the novel object.

### Social Interaction Test (SIT)

Non-sibling, non-cage mates from the same treatment condition, genotype and sex were paired for this task, employing the same arena used for the OFT. Two rats were placed in the arena simultaneously and allowed to interact freely for 10 min. Body-to-body, nose-to-nose, and nose-to-tail contact were tracked automatically using EthoVision XT17 software (Noldus, The Netherlands) to count them as social contact. The total duration of social contact was used as the measure of social interaction. Total social interaction for each pair was averaged, with social interaction analysis conducted on five pairs of rats per group.

### Social Recognition (SR)

SR was assessed using a 3-chamber box, consisting of a Plexiglas box (120×80×40 cm), subdivided into 3 connected identical chambers (40×80 cm). First, the experimental rat was placed into the empty middle chamber and allowed to explore the whole arena for habituation during 10 min. Afterwards, the animal was placed in the waiting cage, the arena cleaned, and an unfamiliar same-sex, younger rat (4 weeks old; stranger 1) was placed inside a plastic grid in the left or right chamber (randomized). An empty grid was present in the other chamber. The experimental rat was placed back in the middle chamber, the doors were opened, and the experimental rat had access to all 3 chambers during a 10 min testing period (sociability trial). Immediately after, the doors were closed, the experimental rat removed, and stranger 1, now the familiar social stimulus, remained in the chamber. The arena was cleaned, and a second unfamiliar same-sex, younger rat (novel social stimulus) was now placed into the previously empty grid. The experimental rat was placed back in the middle chamber, the doors were opened, and the experimental rat explored all chambers for another 10 min (social recognition trial). All trials were recorded with a camera placed above the arena. Exploration of the grids (empty/social stimulus) for both trials was automatically tracked by using EthoVision XT18 software (Noldus, The Netherlands). Exploration of the grid was defined as the nose pointing towards the grid from a distance of 2 cm or less, or directly touching the grid. For the sociability trial, the time spent exploring the grid with the stranger 1 and the empty grid were scored, and the social preference was calculated as:

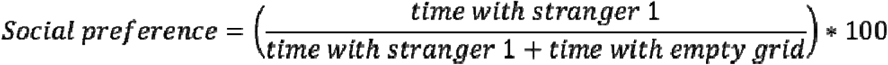

For the social recognition trial, the time spent exploring each grid containing the social stimulus was scored, and the sum of the two corresponded to the total duration of social exploration. The recognition of the novel social stimulus was calculated as:

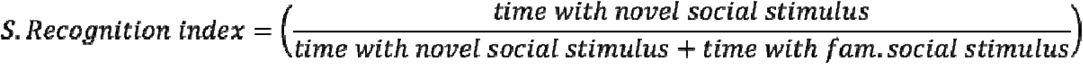

The social recognition index above 0.5 indicated the preference of the novel social stimulus over the familiar (social recognition).

All behavioral tests were conducted in the same experimental room under dim light (approx. 25 Lux), except for the SPT, which was performed under red light in the home cage room. Estrous cycle was measured before starting each test; females in proestrus stage were left in their home cages and tested in one of the following days when they were not in the fertile part of the cycle.

## Notes

### Competing Interest Statement

The authors have declared no competing interest.

